# Time-Resolved Neural Oscillations Across Sleep Stages: Associations with Sleep Quality and Aging

**DOI:** 10.1101/2024.12.16.628645

**Authors:** Antony D Passaro, Alexander Poltorak

**Affiliations:** NeuroLight, LLC 8 Beaver Dam Rd., Pomona, NY 10970

**Author notes:** Corresponding Author: Antony D Passaro, PhD 1209 Lost Creek Blvd Austin, TX 78746.

**Keywords:** EEG analysis, Functional Brain Imaging, Sleep and the Brain, NREM-REM Cycles, Slow Wave Sleep, Aging, Sleep/Wake Physiology

## Abstract

Sleep is a fundamental physiological process critical to cognitive function, memory consolidation, emotional regulation, and overall health. This study investigates the relationship between EEG spectral power dynamics and key sleep metrics, including percentage of N3, biological age, percentage of REM, and total sleep time (TST). Using high-resolution spectral analysis, we examine how power across multiple frequency bands (0.1–50 Hz) evolves temporally across sleep stages and influences sleep architecture. Our results reveal an inverse relationship between high-frequency power (sigma, beta, and gamma) during the N1 and N2 stages and the subsequent percentage of N3, suggesting that excessive low-frequency power in N2 may disrupt the smooth progression into deep sleep. Additionally, we identify a negative correlation between low delta power (0.1–0.5 Hz) during N2 and both percentages of N3 and TST, challenging traditional views on the role of delta activity in sleep regulation. These findings advance the understanding of how brain activity across frequencies modulates sleep depth and duration, with implications for addressing age-related sleep declines.

**Statement of Significance:** This research highlights novel insights into the relationships between EEG spectral power dynamics and sleep architecture, offering a deeper understanding of how brain activity influences critical sleep metrics such as N3 percentage, REM percentage, and total sleep time. By revealing unexpected findings, such as the inverse relationship between low-frequency power during N2 and N3 duration, this study challenges conventional sleep science paradigms. These findings have significant implications for addressing age-related sleep declines and designing brain-computer interfaces (BCIs) to optimize sleep. By targeting specific frequency bands and leveraging real-time feedback, this work paves the way for personalized, non-invasive sleep modulation therapies, revolutionizing clinical and home-based sleep interventions.

## Introduction

Understanding how EEG spectral activity evolves during transitions between wakefulness and successive sleep stages is critical for clarifying the mechanisms that govern sleep architecture. Transitions into NREM and REM sleep are not discrete switches but gradual shifts in oscillatory and aperiodic dynamics, and the spectral composition of these transitions can shape the depth, continuity, and duration of subsequent stages. Across the night, spectral power in specific bands not only defines individual stages but also correlates with global outcomes such as the proportion of N3, the amount of REM, and total sleep time. Characterizing these temporally resolved spectral changes therefore provides a window into the regulatory processes that determine how sleep unfolds and how it supports cognitive and physiological function. Such insights hold translational value for identifying biomarkers of disordered sleep, refining theories of sleep regulation, and informing targeted interventions aimed at improving sleep quality across the lifespan.

### EEG Sleep Dynamics

The transition from wakefulness into sleep is marked by a progressive reorganization of spectral activity rather than a discrete switch in brain state. During sleep onset (Awake→N1), high-frequency beta and gamma activity typically associated with cortical arousal decreases, while alpha rhythms fragment and give way to increasing theta power (Marzano et al., 2013; Lustenberger et al., 2019). These shifts are thought to reflect a gradual disengagement from external processing and a buildup of network synchronization, yet the precise relationship between pre-sleep high-frequency activity and the stability of sleep onset remains incompletely understood.

The N1→N2 transition is characterized by the consolidation of theta rhythms and the appearance of spindles and K-complexes, signaling the stabilization of NREM sleep. While spindles are consistently associated with memory consolidation and sleep protection, their role in shaping subsequent slow-wave activity is debated. Some studies suggest that excessive spindle activity may reduce the depth of following N3, while others indicate a complementary role in facilitating deeper sleep (Fogel & Smith, 2011; Fernandez & Lüthi, 2020). The variability in these findings underscores that the influence of N1 and N2 dynamics on downstream architecture remains unresolved.

The N2→N3 transition involves the progressive amplification of slow oscillations in the delta range. SWA is widely accepted as the electrophysiological hallmark of homeostatic recovery, yet its initiation during this transition is not fully explained by prior wake time alone. For example, very low-frequency power (<0.5 Hz) sometimes appears prematurely in N2, raising the possibility that early emergence of slow activity can interfere with smooth entry into consolidated N3 (Mander et al., 2017; Fattinger et al., 2017). Understanding how sigma, beta, and low-delta power interact in this transitional window remains an open question.

Finally, the N2→REM transition presents one of the most striking reversals in spectral composition: delta and sigma activity diminish while higher-frequency beta/gamma activity re-emerges, resembling wake-like cortical activation (Nir & Tononi, 2010; Siclari et al., 2020). While the ultradian timing of NREM and REM cycles is well established, much less is known about how the spectral profile of N2 influences REM onset and duration. Some evidence suggests that excessive slow-wave buildup during N2 can delay REM entry (Bódizs et al., 2005), but systematic transition-level analyses have been scarce.

Taken together, prior work has defined many of the spectral features associated with canonical stages, but the precise ways in which **transitions themselves** shape the depth and continuity of subsequent stages remain poorly characterized. This gap motivates a closer look at temporally resolved spectral dynamics across transitions, which may provide more direct insight into the regulatory mechanisms of sleep than stage-averaged measures alone.

### N3 Sleep

N3 sleep, characterized by high-amplitude slow waves in the delta range (0.5–4 Hz), is a key marker of restorative sleep and an index of homeostatic recovery. High levels of N3 are consistently associated with better cognitive performance, emotional regulation, and metabolic health, while reduced N3 is linked to memory deficits, mood disturbances, and increased risk for neurodegenerative disease (Mander, Winer, & Walker 2017; Varga et al. 2016). Demographically, N3 declines sharply with age and is typically reduced in males compared to females, suggesting that both age and sex significantly shape the distribution of slow-wave sleep (Carrier et al. 2011; Redline et al. 2004). Interindividual variability in N3 also predicts susceptibility to sleep loss, with those exhibiting lower baseline N3 showing greater cognitive impairment after deprivation (Rupp et al. 2012).

Recent studies have demonstrated that both high and low proportions of N3 carry distinct implications. Elevated N3 may reflect robust homeostatic processes and is often linked to improved memory consolidation (Diekelmann & Born 2010), but unusually high N3 can also signal compensatory rebound following sleep loss or fragmented sleep, and may disrupt the ultradian balance between NREM and REM cycles (Lecci et al. 2017). Conversely, reduced N3 is associated not only with aging but also with psychiatric and metabolic disorders, including depression, type 2 diabetes, and hypertension (Fang et al. 2015; Kim et al. 2013). Importantly, the proportion of N3 often predicts subsequent REM duration, supporting the view that deep NREM recovery processes set the stage for REM expression (Dijk 2009; Bódizs et al. 2005).

Beyond delta activity, additional frequency bands modulate N3 expression. Spindle activity (sigma band, 11–15 Hz) in N2 has been associated with the build-up of subsequent slow-wave activity, while elevated beta power during NREM predicts reduced N3% and greater vulnerability to arousals (Fernandez & Lüthi 2020; Helfrich et al. 2018). These findings highlight that the depth and continuity of N3 are not determined by delta oscillations alone but emerge from an interaction of multiple spectral components across transitions. Despite these insights, the specific frequency signatures that promote or disrupt sustained N3 remain incompletely understood, underscoring the need for transition-focused analyses.

### REM Sleep

REM sleep expression depends not only on circadian timing but also on the spectral and structural features of preceding NREM. Longer or deeper NREM bouts predict more robust REM episodes, while insufficient slow-wave buildup can delay or suppress REM onset (Dijk 2009; Bódizs et al. 2005). This stage-to-stage dependency highlights that REM is not an isolated process but emerges from the evolving dynamics of prior NREM cycles.

REM percentage (REM%) relative to total sleep time is a widely used index of sleep quality. Reductions in REM% are consistently associated with depression, anxiety, and post-traumatic stress disorder, while higher REM% correlates with enhanced emotional regulation and cognitive flexibility (Cartwright et al. 1998; Cai et al. 2009; Goldstein & Walker 2014). In neurodegenerative disease, especially Parkinson’s and dementia with Lewy bodies, REM disruptions are an early hallmark, often presenting as REM behavior disorder and reduced atonia (Arnulf 2012; Boeve et al. 2007). Recent work also suggests that diminished REM% is linked to aging-related impairments in emotional memory consolidation (Rosales-Lagarde et al. 2012; Mander et al. 2017).

Despite these associations, the spectral determinants of REM% remain poorly understood. N2 activity in sigma and delta bands has been hypothesized to prime REM expression, while higher beta/gamma activity may facilitate the wake-like cortical activation characteristic of REM. Yet systematic analyses of how NREM spectral composition predicts REM quantity and quality are scarce. Identifying these transition-level spectral predictors of REM is therefore critical for clarifying the mechanisms that govern emotional regulation, memory consolidation, and vulnerability to disease across the lifespan.

### Sleep Duration and Spectral Power

Total sleep time (TST) and sleep continuity remain among the most reliable markers of sleep health, with reductions linked to cognitive decline, mood disturbances, and metabolic dysfunction (Lo et al. 2016; Medic et al. 2017). Recent spectral studies suggest that both oscillatory and aperiodic activity play important roles in shaping sleep duration. Greater delta and slow oscillatory power before and during early NREM sleep is associated with longer and more consolidated TST, consistent with the role of slow waves in dissipating homeostatic pressure (Mander et al. 2017; Djonlagic et al. 2021). In contrast, elevated beta and gamma activity during NREM predicts fragmentation and shorter duration, often reflecting hyperarousal or sleep pathology (Kremen et al. 2019; Helfrich et al. 2021).

Importantly, interindividual differences in baseline spectral profiles partly explain trait-like variability in sleep need. Individuals with naturally higher SWA tend to sustain longer TST and show greater resilience to sleep restriction, whereas those with reduced SWA exhibit increased vulnerability to cognitive deficits after sleep loss (Rupp et al. 2012; Lecci et al. 2017). Beyond oscillatory power, flattening of the aperiodic slope has been associated with lighter, more fragmented sleep, suggesting that non-oscillatory components of the EEG may also serve as markers of sleep continuity (Lendner et al. 2020). Despite these advances, the precise ways in which spectral dynamics during stage transitions predict TST and efficiency remain largely unexplored. Clarifying how transition-level spectral features shape continuity offers a path toward more mechanistic markers of sleep health across individuals and conditions.

### Age-Related Changes in Sleep Architecture

Aging is consistently associated with shorter total sleep time, reduced N3 and REM sleep, and greater fragmentation, with increased transitions to lighter stages and more frequent awakenings (Mander, Winer, & Walker 2017; Scullin & Bliwise 2015). Across the lifespan, slow-wave activity (SWA) shows the steepest decline, decreasing by nearly 40–50% between young adulthood and middle age and continuing to erode into older adulthood (Muehlroth & Werkle-Bergner 2020; Helfrich et al. 2018). This reduction in SWA has been linked to impairments in memory consolidation, executive function, and vulnerability to neurodegenerative disease, particularly Alzheimer’s disease (Mander et al. 2017; Lucey et al. 2019).

Spectral changes with age extend beyond delta power. Older adults show reduced spindle density and altered sigma power, which are thought to impair the ability of N2/N3 to stabilize sleep and support memory processes (Fogel et al. 2017; Muehlroth et al. 2019). High-frequency alpha and beta intrusions increase with age, correlating with lighter, more fragmented sleep and reduced subjective sleep quality (Helfrich et al. 2018). Recent evidence also points to flattening of the aperiodic 1/f slope as a marker of age-related cortical changes in sleep physiology (Lendner et al. 2020).

Despite these advances, key uncertainties remain. It is not well understood how age-related alterations in spectral dynamics at transitions — for example, from N2 into N3 or from N2 into REM — constrain sleep continuity. Moreover, while reduced SWA is robustly linked to aging, interindividual variability is large: some older adults maintain relatively preserved slow-wave activity and exhibit resilience in memory and cognitive function (Muehlroth & Werkle-Bergner 2020). Identifying the spectral features at stage transitions that predict preserved versus disrupted sleep in aging remains an open question and a critical step for understanding vulnerability to cognitive decline.

### The Current Study

Despite decades of work on EEG spectral power and sleep architecture, key questions remain unresolved. Most prior research has averaged spectral activity across entire stages, overlooking the temporally precise dynamics that occur at transitions such as Awake→N1, N1→N2, N2→N3, and N2→REM. While delta activity has been extensively studied as a marker of sleep depth, the contributions of other frequency bands—including sigma, beta, and gamma—as well as aperiodic activity remain poorly defined. Moreover, it is not well understood how spectral features at these transitions shape global outcomes such as N3%, REM%, and total sleep time, or how these relationships vary across the lifespan.

This study addresses these gaps by examining spectral power with fine temporal resolution during stage transitions and testing how these dynamics predict key metrics of sleep depth, continuity, and architecture. By linking transition-level spectral changes to N3%, REM%, TST, and age, we aim to refine understanding of the mechanisms that regulate sleep organization and identify spectral markers of vulnerability to fragmentation and aging. These insights may ultimately inform the development of biomarkers and interventions designed to preserve or enhance sleep quality.

## Methods

### Participants and Data Collection

EEG and polysomnographic (PSG) data were obtained from the Stanford Technology Analytics and Genomics in Sleep (STAGES) study (Zhang et al., 2018), which included 1137 subjects (mean age 46.35 ± 14.05 years). EEG recordings were sampled at 200 Hz, 250 Hz, or 500 Hz, depending on the subject, and were subsequently down-sampled to 100 Hz to ensure uniformity in the analysis. PSG recordings included standard EEG channels (e.g., F3, F4, C3, C4, O1, O2) for the detection of sleep stages and other relevant physiological measurements.

### Sleep Stage Transition Selection

To investigate sleep stage transitions, only subjects who demonstrated at least 5 minutes of contiguous data within a sleep stage followed by a transition to the next sleep stage with a minimum of 5 more minutes were included in the analysis. This ensured that each stage transition contained sufficient data for spectral analysis. Specifically, four transitions were analyzed:

- Awake to N1 (n = 425)
- N1 to N2 (n = 153)
- N2 to N3 (n = 492)
- N2 to REM (n = 538)

For each transition, data from the five minutes preceding and the five minutes following the transition were extracted for further analysis. This resulted in 10-minute segments (6000 time points) per subject and transition, which were then used for spectral analysis.

### Time-Frequency Analysis

EEG data were segmented into continuous 10-minute windows centered on each transition and analyzed at high temporal resolution. Signals were transformed into the time–frequency domain using a multitaper approach with discrete prolate spheroidal sequence (DPSS) tapers, which provide stable spectral estimates while minimizing leakage across neighboring frequencies. Because electrode configurations varied across sites in this dataset, we restricted analyses to C3 and C4, which were the universally available central derivations. These electrodes are standard for staging in polysomnography and provided reliable spectral estimates of both slow-wave and spindle activity. Power from C3 and C4 was averaged to yield a single measure per participant. Both the spectral smoothing and the temporal windows were scaled systematically with frequency:

- Spectral tapers: The number and bandwidth of DPSS tapers were scaled *logarithmically with frequency*. At higher frequencies (beta/gamma), broader bandwidths stabilized power estimates, while at lower frequencies, narrower bandwidths preserved specificity of slow oscillations.
- Temporal windows: The length of the analysis window was scaled as the inverse of the natural logarithm of frequency (1/ln[f]). Very low frequencies (<1 Hz) were estimated using long windows spanning multiple seconds, ensuring multiple oscillatory cycles were included, while higher frequencies were estimated with progressively shorter windows to maximize temporal precision.

Spectral power was estimated continuously in 100 ms steps across the 10-minute windows, yielding a dense time–frequency representation. Output frequencies ranged from 0.1 to 50 Hz and were interpolated onto a linear frequency axis for interpretability. Within each subject, power was normalized relative to the mean across the transition window to reduce baseline variability.

This procedure ensured that slow and fast oscillations were estimated with appropriate resolution, minimized leakage, and provided a continuous map of spectral dynamics during stage transitions.

### Linear Regression Analysis for Sleep Stage Transitions

To examine linear trends in spectral power during each transition, linear regressions were performed at the individual-subject level. For each subject, spectral power at each frequency (0.1–50 Hz) was regressed against time within the 10-minute segments surrounding each stage transition. This was done separately for each transition type (Awake→N1, N1→N2, N2→N3, N2→REM). For each regression, the slope of the regression line was extracted for every frequency.

At the group level, subject-level slopes were entered into one-sample *t*-tests against zero to assess whether the average slope differed significantly across participants. This procedure treated the subject as the unit of analysis and avoided inflating degrees of freedom by treating multiple epochs as independent. Group-level *p*-values were corrected for multiple comparisons across frequencies (see below). Significant effects are reported where the group distribution of slopes showed consistent increases or decreases in power across subjects.

### Overnight Regression Analysis of Spectral Power

To characterize spectral changes across the entire night, we conducted regressions at the **subject level** within each sleep stage (Awake, N1, N2, N3, REM). For each subject, spectral power at each frequency (0.1–50 Hz) was regressed against time within that stage, yielding one slope estimate per frequency per subject. Group-level inference was then performed by testing these subject-level slopes against zero across participants, using one-sample *t*-tests. Results were plotted as the mean group-level slope ± SEM as a function of frequency for each stage, with false discovery rate (FDR) correction applied across the full frequency range.

### Correlation Analysis: Spectral Power and Sleep Metrics

To explore the relationship between spectral power and key sleep-related metrics, a series of Pearson correlation analyses were performed. Four metrics were considered:

1. Percentage of N3 sleep relative to total sleep time (N3%)
2. Percentage of REM sleep relative to total sleep time (REM%)
3. Total sleep time (TST)
4. Age of the subject

For each subject, the spectral power calculated for all 10-second epochs within each sleep stage (awake, N1, N2, N3, REM) was averaged to yield a mean power value for each frequency (0.1 Hz to 50 Hz) within each stage. These average spectral power values were then correlated with each of the four metrics across all subjects. The FDR correction was applied to account for multiple comparisons (p = 0.125 due to 4 sets of comparisons), and significant correlations were identified and reported.

### Correlation Analysis by Sleep Stage

- Awake: Correlations between spectral power during the awake period and the four metrics (N3%, REM%, TST, and age) were calculated to assess how pre-sleep brain activity might influence sleep architecture and quality.
- N1: Correlations were computed to investigate how early-stage sleep (N1) activity relates to sleep outcomes and age.
- N2 and N3: N2 and N3 correlations were critical for understanding the relationship between deeper NREM sleep activity (e.g., delta waves) and metrics such as N3% and age.
- REM: The correlation between REM spectral power and each metric provided insight into how REM activity is linked to sleep health and age.

### Correlation Analysis: Temporally Regressed T-Values and Sleep Metrics

In addition to correlating raw spectral power, a secondary analysis involved correlating the temporally regressed t-values from the linear regression analyses described above with the four sleep metrics (N3%, REM%, TST, and age). This allowed for the examination of how linear changes in spectral power over time during each stage and frequency band were associated with sleep quality and aging.

### Correlation Analysis by Sleep Stage

- Awake: Temporal changes in spectral power during the awake period were correlated with sleep metrics, providing insight into how pre-sleep changes in brain activity might predict sleep quality (e.g., more REM or longer TST).
- N1: Temporal trends in N1 activity were examined for their association with sleep metrics, such as how gradual changes in early-stage sleep might correlate with increased N3% or TST.
- N2 and N3: The temporal evolution of spectral power in N2 and N3 stages was particularly relevant for understanding slow-wave activity changes over the course of the night and how these changes correlate with sleep metrics such as N3% and TST.
- REM: Temporal changes in REM spectral power were examined to assess whether gradual increases or decreases in activity over the night were associated with changes in REM%, TST, or age.

The FDR correction was applied to both the spectral power correlations and temporally regressed t-value correlations, and statistically significant relationships were highlighted. These analyses provided insight into how both static and dynamic changes in spectral power during sleep correlate with overall sleep quality and age.

### Correction for Multiple Comparisons

Spectral estimates were computed separately for each frequency, producing a large number of statistical tests. To control the false discovery rate, we applied Benjamini–Hochberg FDR correction (Benjamini & Hochberg, 1995) across all frequencies tested within each analysis. Specifically:

- For transition regressions, all frequencies from 0.1–50 Hz were corrected together within each transition type (Awake→N1, N1→N2, N2→N3, N2→REM).
- For overnight regressions, all frequencies were corrected together within each sleep stage (Awake, N1, N2, N3, REM).
- For correlations with sleep metrics (N3%, REM%, TST), corrections were applied across all tested frequencies simultaneously.

In total, *N* = 500 frequency bins were included per analysis, and reported *p*-values therefore reflect effects that survive correction across the full frequency spectrum.

All analyses were correlational and exploratory in nature. Regression slopes and associations between spectral power and sleep metrics (N3%, REM%, TST) were used to characterize statistical relationships but should not be interpreted as evidence of causality. No manipulations were performed, and the analyses were designed to describe associations rather than infer mechanistic directionality.

## Results

### Time-Frequency Analysis of Sleep Stage Transitions

#### Awake to N1 Transition

During the transition from wakefulness to N1 (**Figure 1**), spectral power decreased significantly in the beta (15–30 Hz) and gamma (30–50 Hz) ranges, while power increased in delta (0.5–4 Hz) and theta (4–7 Hz). A modest increase was also observed in the sigma band (12–15 Hz). These results indicate broad reductions in high-frequency activity and concomitant increases in lower-frequency power at the onset of sleep.

**Figure 1.**
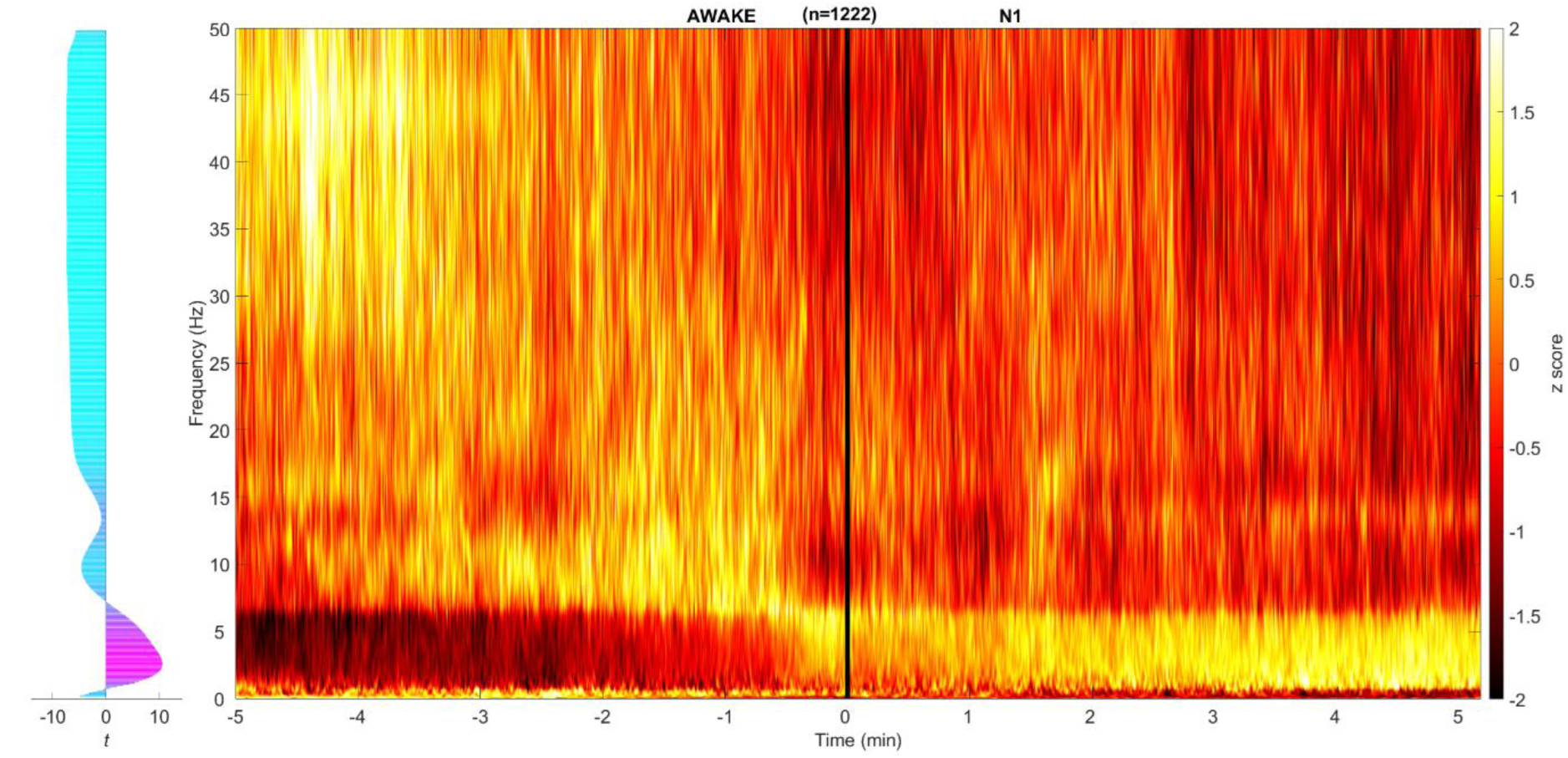
Spectral Power Changes During the Transition from Wakefulness to N1 Sleep. Time-frequency analysis of EEG spectral power changes during the transition from wakefulness to N1 sleep. The heatmap represents the averaged power spectra across all participants, with frequency on the y-axis (0.1 to 50 Hz) and time relative to the transition (in minutes) on the x-axis. Lighter colors, such as orange and yellow to white, indicate increased power, while darker colors, such as red and dark red to black, represent decreases. *t* values corresponding to temporal regressions over the 10 min time period are plotted to the left with increasing *t* values as power increases over time displayed as purple to magenta and decreasing *t* values as power decreases over time displayed as blue to cyan.

#### N1 to N2 Transition

The transition from N1 to N2 (**Figure 2**) was characterized by further decreases in beta and gamma power. Delta and theta power continued to increase, and sigma power showed a more pronounced elevation relative to the preceding transition.

**Figure 2.**
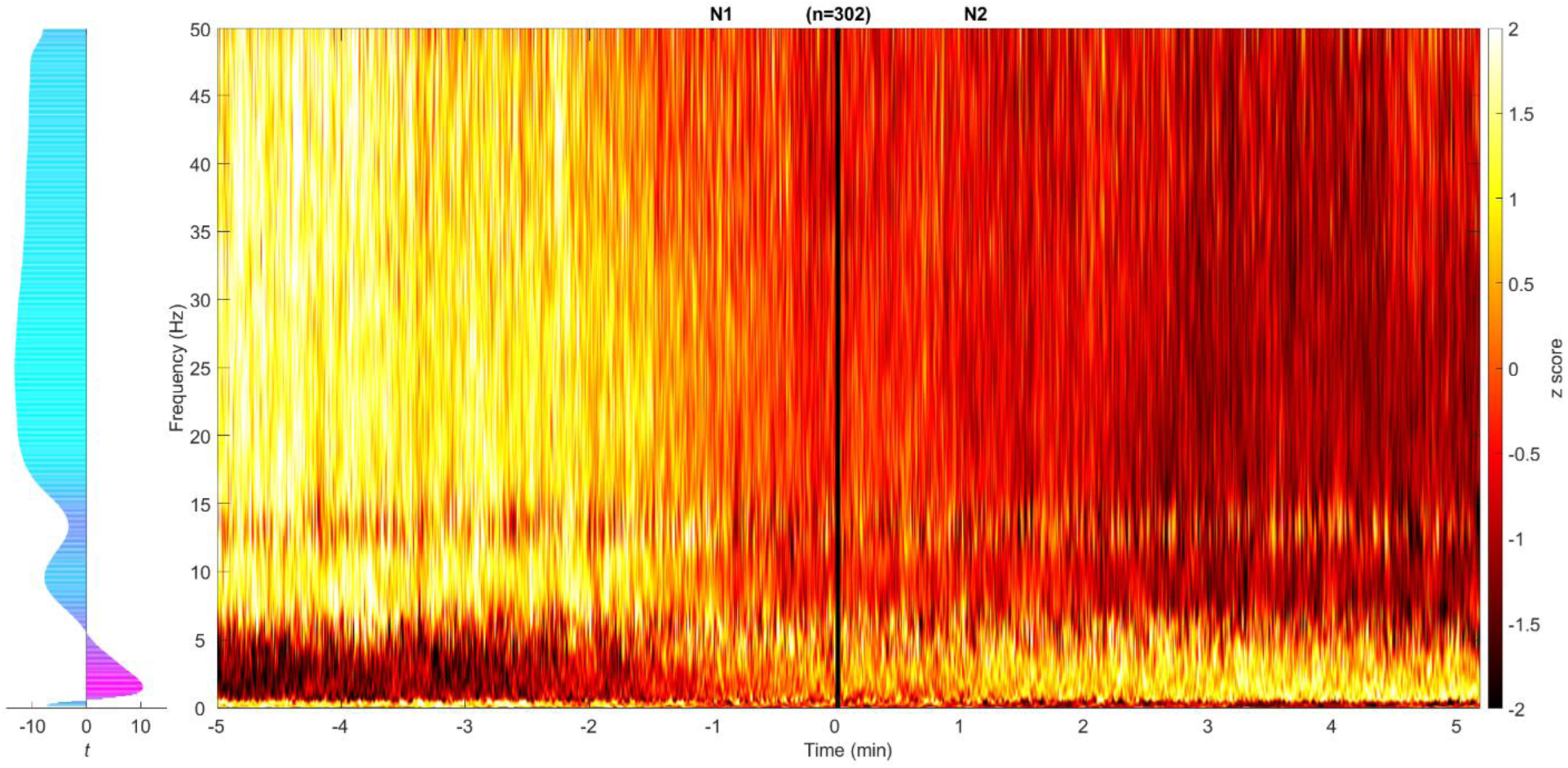
Spectral Power Changes During the Transition from N1 to N2 Sleep. Time-frequency analysis of EEG power changes during the transition from N1 to N2 sleep for a −5 to 5 min period with corresponding temporal regressions in power plotted on the left.

#### N2 to N3 Transition

The transition from N2 to N3 (**Figure 3**) showed fewer significant changes compared to earlier transitions. Power decreases were observed across alpha (8–12 Hz), beta, and gamma bands, while delta power (0.5–4 Hz) increased, consistent with the onset of slow-wave activity.

**Figure 3.**
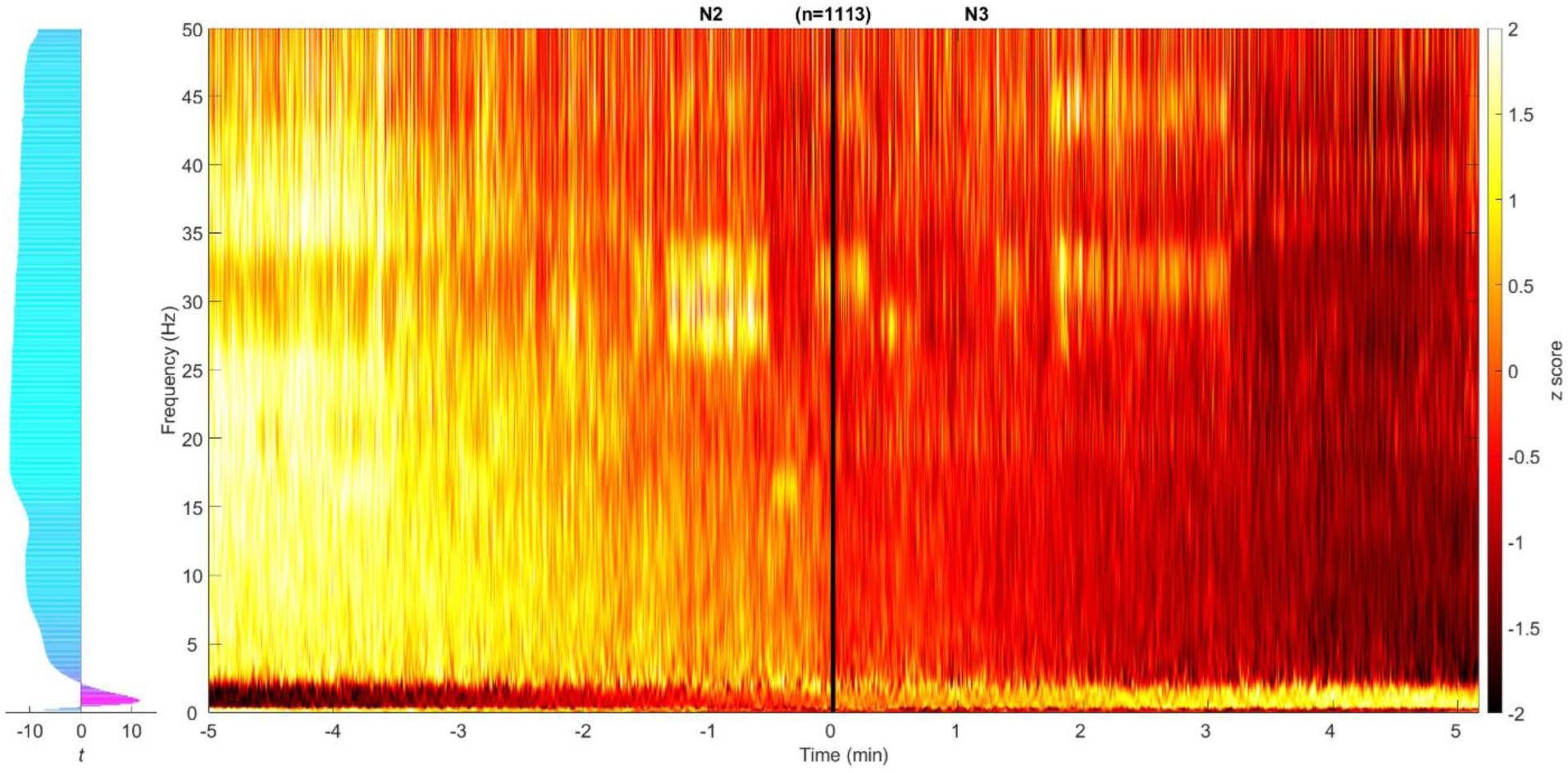
Spectral Power Changes During the Transition from N2 to N3 Sleep. Time-frequency analysis of EEG power during the transition from N2 to N3 sleep for a −5 to 5 min period with corresponding temporal regressions in power plotted on the left.

#### N2 to REM Transition

During the N2 to REM transition (**Figure 4**), spectral power increased in beta and gamma frequencies, while decreases were observed in delta and sigma power. These changes reflect the mixed-frequency profile of REM sleep relative to N2.

**Figure 4.**
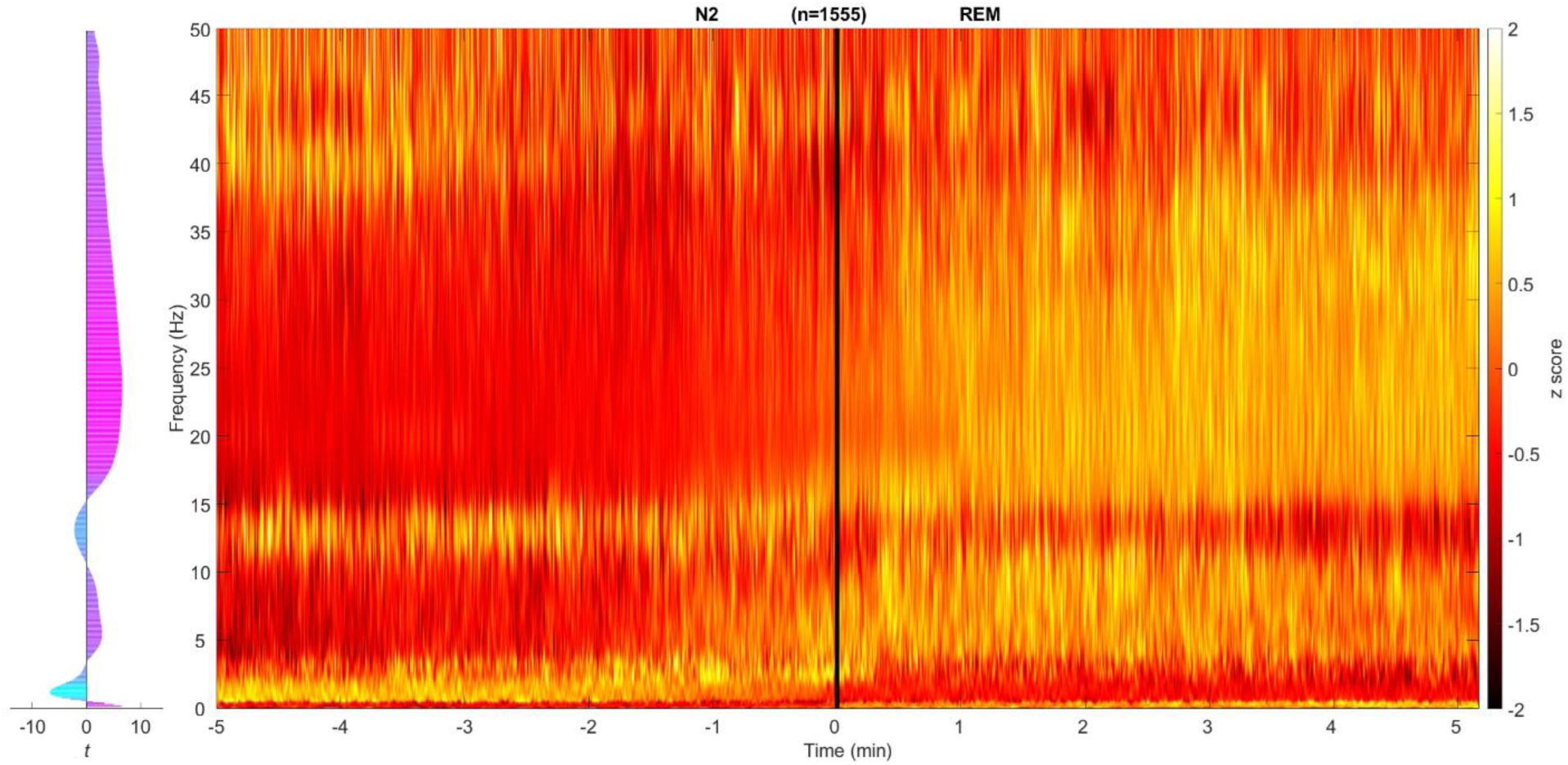
Spectral Power Changes During the Transition from N2 to REM Sleep. This figure illustrates the time-frequency analysis of EEG power changes during the transition from N2 to REM sleep for a −5 to 5 min period with corresponding temporal regressions in power plotted on the left.

### Linear Regression Analysis for Sleep Stage Transitions

We tested whether spectral power changed systematically across the night within each stage (**Figure 5**). Fluctuations were observed in all five stages, though only a subset reached statistical significance after correction. In N2, delta power (0.5–4 Hz) decreased significantly across the night. This effect was strongest in the lower delta range (0.5–2 Hz), while higher delta frequencies (2–4 Hz) showed a weaker trend. No consistent changes were observed in sigma, beta, or gamma power during N2. In N3, infraslow activity (<0.5 Hz) increased across the night, in contrast to the decrease observed in higher delta frequencies. This divergence between infraslow and delta bands was consistent across participants and suggests different temporal trajectories for very slow oscillations versus conventional slow-wave activity. No other bands in N3 exhibited significant trends. In REM, modest decreases were observed in sigma and low beta activity (12–20 Hz), though these did not survive correction across all frequencies. N1 and wakefulness did not show consistent linear changes. These analyses indicate that changes in spectral power across the night are largely confined to the delta and infraslow ranges, and that the direction of change differs by stage.

**Figure 5.**
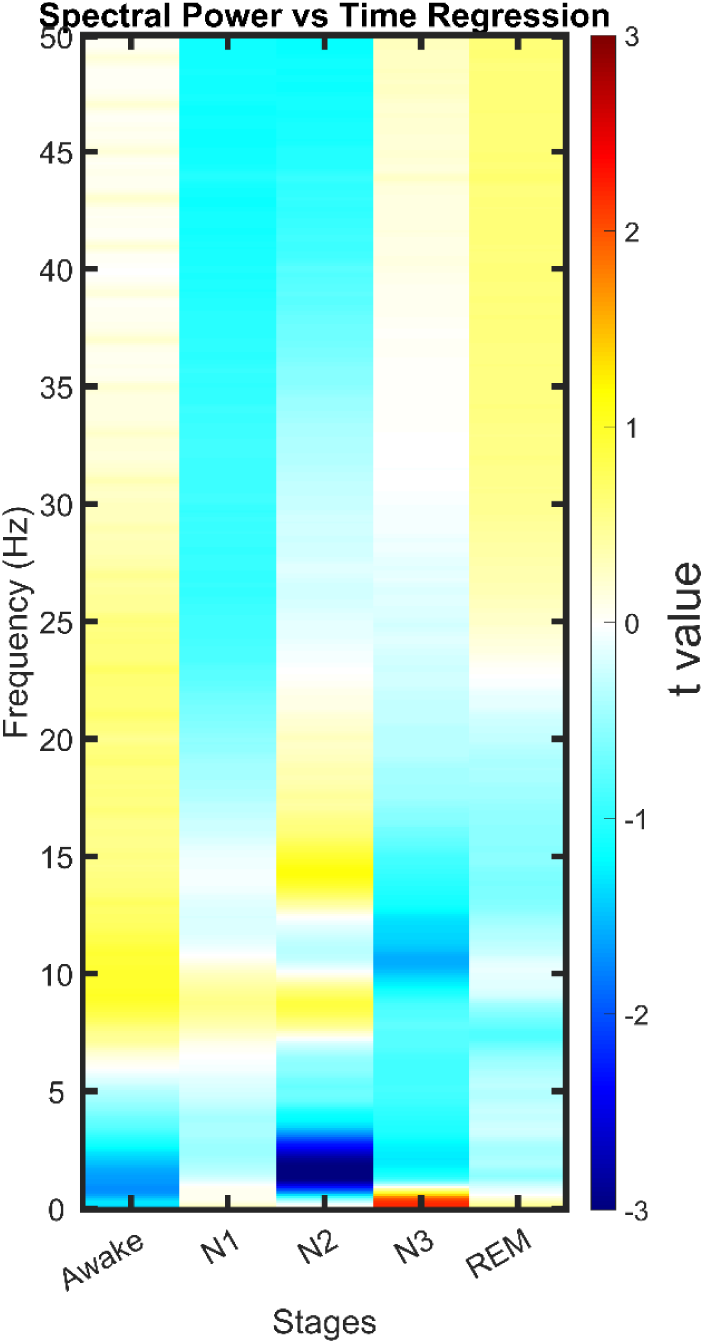
Temporal regression of spectral power across sleep stages and frequencies. This figure shows the results of linear regression between time and spectral power across different sleep stages (awake, N1, N2, N3, and REM) and frequencies from 0.1 to 50 Hz. Warm colors (orange to red) represent positive regressions, indicating increases in spectral power over time, while cool colors (cyan to blue) represent negative regressions, indicating decreases. Statistically significant regions (p < 0.05, FDR-corrected) are colored.

### Correlation Analysis of Spectral Power and Sleep Metrics

#### N3 Percentage

Several frequency bands showed significant associations with the proportion of time spent in N3 (**Figure 6**). Sigma power (12–15 Hz) during N1 was negatively correlated with N3%, as was infraslow activity (<0.5 Hz) during N2. Additional negative correlations were observed between N3% and beta/gamma power during both N2 and REM. Together, these results suggest that higher spindle- and high-frequency activity in lighter stages is consistently associated with reduced time in N3.

**Figure 6.**
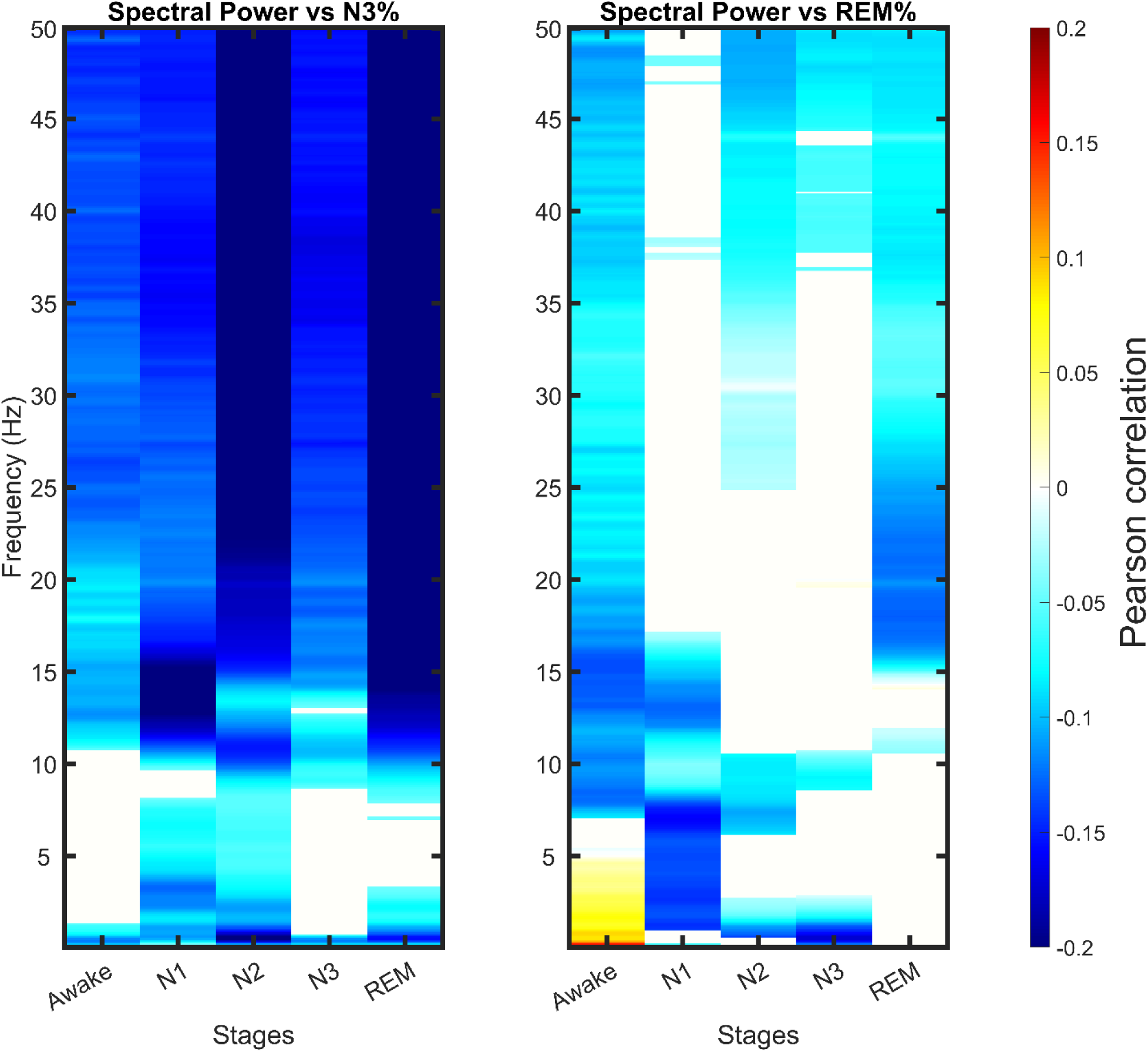
Correlations of spectral power with N3 percentage (left) and REM percentage (right) across sleep stages and frequencies. This figure displays the correlation results between spectral power and the percentage of N3 sleep (left) and REM sleep (right) for each sleep stage (awake, N1, N2, N3, and REM) and frequency from 0.1 to 50 Hz. Warm colors (yellow to red) represent positive correlations, where higher spectral power is associated with greater N3 or REM percentage, while cool colors (cyan to blue represent negative correlations. Statistically significant regions (p < 0.05, FDR-corrected) are colored.

#### REM Percentage

REM% was inversely related to higher-frequency activity across multiple stages. Specifically, alpha through gamma power (8–50 Hz) during N1, N2, and N3 correlated negatively with REM%. Conversely, delta activity (0.5–4 Hz) during wakefulness correlated positively with REM%, suggesting that pre-sleep slow activity is associated with greater REM later in the night.

#### Sleep Duration

Total sleep time (TST) was also linked to spectral activity (**Figure 7**). High-frequency power (alpha through gamma, 8–50 Hz) during N2, N3, and REM correlated negatively with TST. In contrast, delta power (0.5–4 Hz) during wakefulness was positively correlated with TST. These findings indicate that low pre-sleep delta power is associated with shorter sleep durations, while high cortical activation during sleep (alpha–gamma) predicts reduced sleep continuity

**Figure 7.**
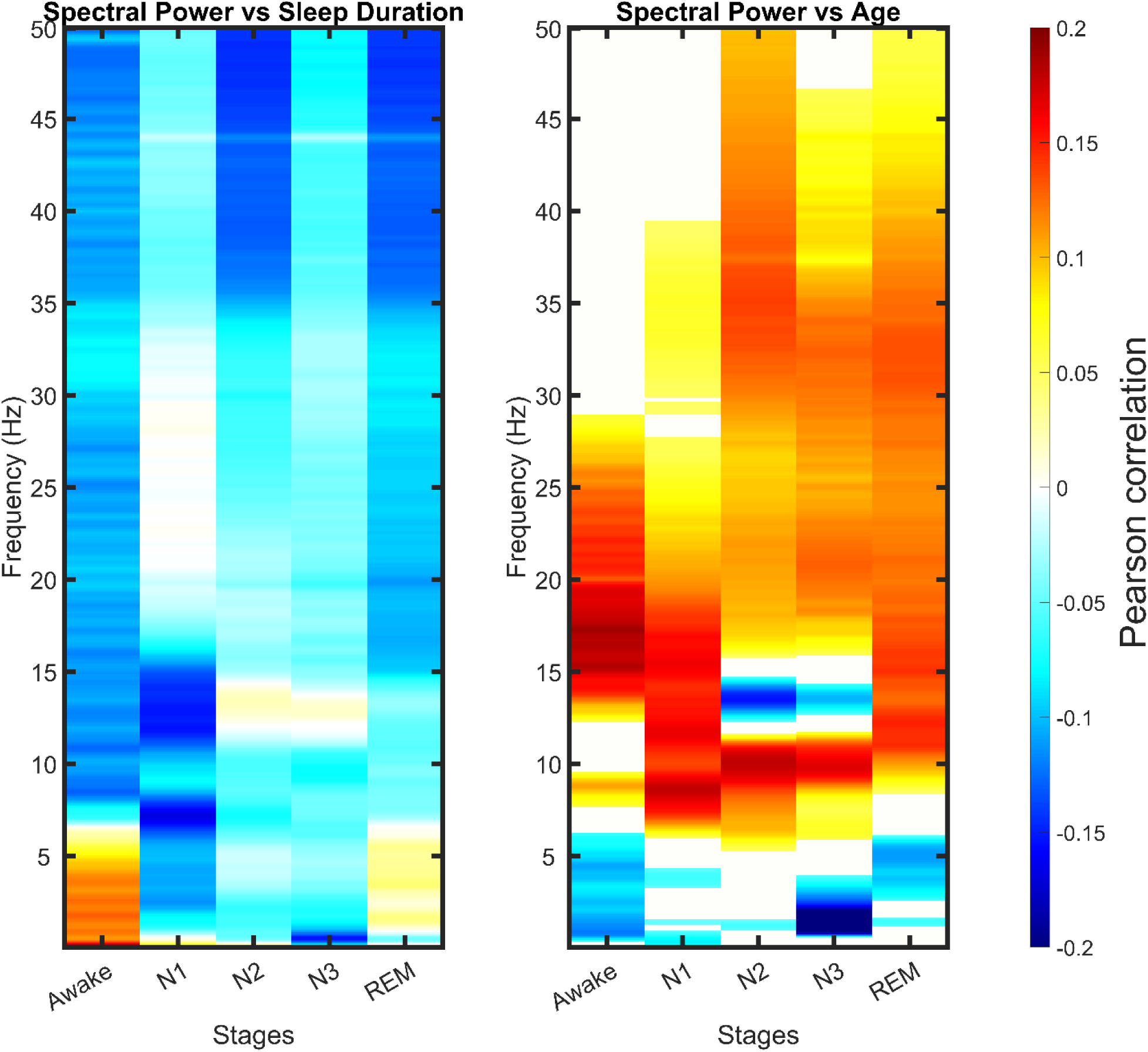
Correlations of spectral power with sleep duration (left) and age (right) across sleep stages and frequencies. This figure presents the correlation results between spectral power and total sleep duration (left) and age (right) for each sleep stage (awake, N1, N2, N3, and REM) and frequency range from 0.1 to 50 Hz. Warm colors (yellow to red) indicate positive correlations, showing that higher spectral power is associated with longer sleep duration or greater age, while cool colors (cyan to blue) indicate negative correlations. Statistically significant regions (p < 0.05, FDR-corrected) are colored.

#### Age

Significant associations with age were observed across multiple stages (Figure 7). During wakefulness, beta and gamma power correlated positively with age, while delta power showed a modest negative correlation. Across N1, N2, and REM, alpha and beta power were positively correlated with age. Within N3, age was negatively correlated with both delta and sigma power. These findings replicate well-established reductions in slow-wave and spindle activity with aging, alongside relative increases in higher-frequency activity.

### Correlation Analysis of Temporally Regressed T-Values and Sleep Metrics

#### N3 Percentage

Greater decreases in sigma and low beta power (12–19 Hz) during wakefulness were associated with higher N3% across the night (**Figure 8**). Within N2, increases in delta power (0.5–4 Hz) over time correlated positively with N3%. In N3 itself, two distinct patterns emerged: infraslow power (<0.5 Hz) increases were positively correlated with N3%, while decreases in higher delta to theta power (1–7 Hz) were also associated with greater N3%. Together, these findings indicate that both pre-sleep activity and within-stage dynamics contribute to the amount of slow-wave sleep obtained.

**Figure 8.**
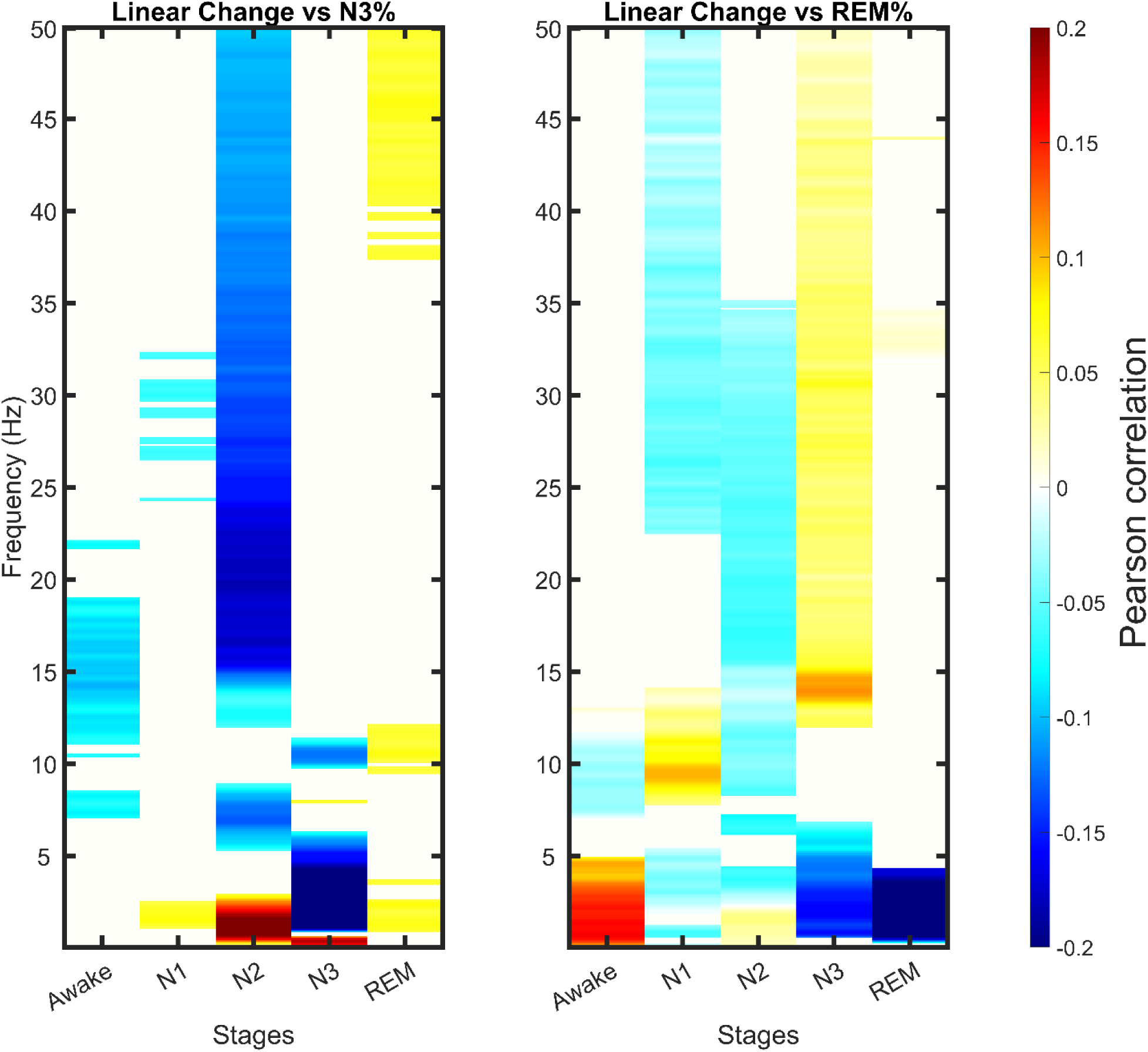
Correlation of temporally regressed spectral power with N3 percentage (left) and REM percentage (right) across sleep stages and frequencies. This figure shows the results of correlations between temporally regressed spectral power and the percentage of N3 sleep (left) and REM sleep (right) for each sleep stage (awake, N1, N2, N3, and REM) and frequency range from 0.1 to 50 Hz. Warm colors (yellow to red) represent positive correlations, indicating that increases in spectral power over time are associated with higher N3 or REM percentages, while cool colors (cyan to blue) represent negative correlations. Statistically significant areas (p < 0.05, FDR-corrected) are colored.

#### REM Percentage

REM% was positively correlated with increases in delta power during wakefulness (**Figure 8**). Within N3, decreases in delta power (0.5–4 Hz) were associated with lower REM%, whereas increases in sigma power (12–15 Hz) were positively associated with REM%. In REM itself, decreases in delta activity correlated negatively with REM%. These findings highlight multiple frequency-specific dynamics that distinguish individuals who obtain greater amounts of REM sleep.

#### Sleep Duration

Total sleep time (TST) was strongly related to pre-sleep dynamics (**Figure 9**). Increases in delta power during wakefulness were positively correlated with TST. Additional positive associations were observed for alpha activity (8–12 Hz) during N1, sigma-to-beta activity (12–25 Hz) during N2, and beta-to-gamma activity (20–40 Hz) during N3. In contrast, decreases in delta power during N2 and N3 were negatively correlated with TST, indicating that sustained slow-wave activity in these stages was linked to shorter total sleep duration.

**Figure 9.**
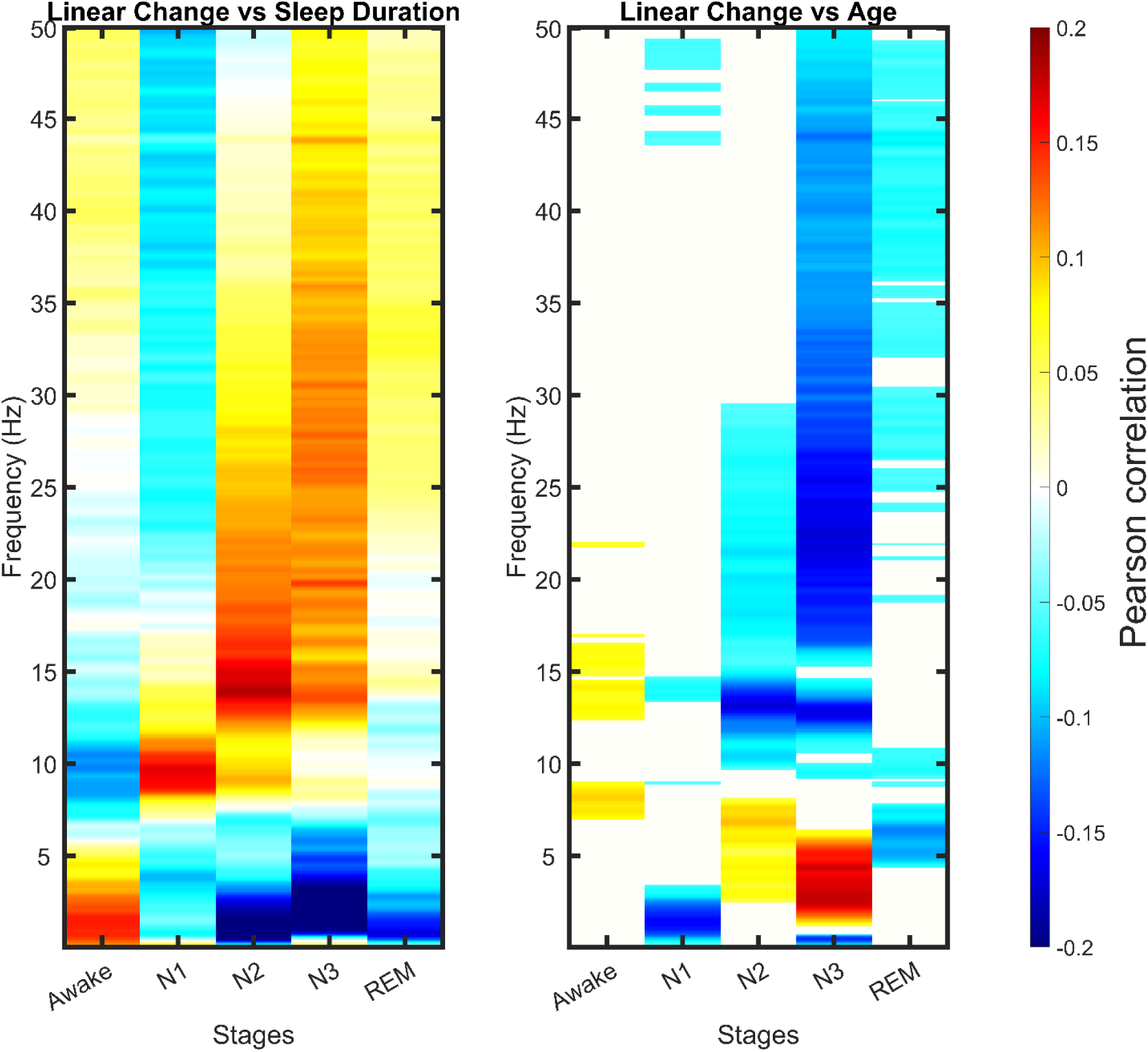
Correlation of temporally regressed spectral power with sleep duration (left) and age (right) across sleep stages and frequencies. This figure illustrates the correlations between temporally regressed spectral power and total sleep duration (left) and age (right) for each sleep stage (awake, N1, N2, N3, and REM) and frequency range from 0.1 to 50 Hz. Warm colors (yellow to red) indicate positive correlations, showing that increases in spectral power over time are associated with longer sleep duration or greater age, while cool colors (cyan to blue) indicate negative correlations. Statistically significant regions (p < 0.05, FDR-corrected) are colored.

#### Age

Temporal regression analyses also revealed age-related associations (**Figure 9**). In N3, older adults exhibited decreases in infraslow activity (<0.5 Hz) over time, coupled with relative increases in higher delta and theta power (1–7 Hz). In REM, age was negatively correlated with decreases in theta activity. These results are consistent with broad age-related shifts in sleep spectral composition across multiple stages.

## Discussion

This study provides a comprehensive analysis of EEG spectral power across different sleep stages and transitions therebetween, confirming established findings while offering novel insights that challenge current literature on sleep. By examining the relationships between EEG spectral power and key metrics—such as N3 percentage, REM percentage, total sleep duration, and age—both in static and temporally regressed forms, we have gained new perspectives on how sleep evolves over time and how brain activity relates to sleep quality and aging. The results highlight several important implications for the regulation of sleep, novel observations that contrast with previous findings, and potential applications for personalized sleep interventions, including brain-computer interfaces (BCIs).

Many of our results corroborate existing knowledge of sleep neurophysiology, particularly regarding transitions from wakefulness into NREM and REM sleep. The decrease in beta and gamma power (16–50 Hz) observed during the transition from wakefulness to N1 and N2 sleep stages reflects the brain’s gradual disengagement from sensory input and cognitive processing as sleep deepens. This finding is consistent with several studies that have documented reductions in high-frequency activity during early sleep as an indicator of reduced cortical arousal (Cantero et al. 2004); (Cantero et al., 2004; Gais et al., 2007)(Iber et al. 2007).

Similarly, the observed increase in lower-frequency power—particularly in the delta (0.1–4 Hz) and theta (4–7 Hz) bands—as subjects transition from wakefulness into NREM stages, which mirrors the well-established pattern of increasing cortical synchronization in the early stages of sleep (Steriade, McCormick, and Sejnowski 1993)( Borbély & Achermann, 1999). These low-frequency oscillations are known to play a crucial role in early sleep stages, reflecting the brain’s shift into a restorative state (D.-J. Dijk 2009).

Our findings also reaffirm the role of sigma power (12–15 Hz) in sleep spindle activity during N2 sleep, which is associated with sleep stability, memory consolidation, and cognitive processing (Fogel and Smith 2011); (Mölle et al. 2002)(. Consistent with earlier research, we observed that sleep spindle activity decreases during transitions from N2 to N3 (Figure 3), which reflects the reduction of spindle generation as the brain enters deep NREM sleep, dominated by slow-wave activity (SWA) (Andrillon et al. 2011).

The increase in delta power during N3 sleep, particularly in the very low delta range (0.1–0.4 Hz), further corroborates the established understanding of slow-wave sleep (SWS) as the most restorative stage of sleep. This finding aligns with the extensive literature demonstrating that slow-wave activity (SWA) is critical for neural recovery, memory consolidation, and overall brain plasticity during sleep (Tononi and Cirelli 2006); (Diekelmann and Born 2010). Additionally, the increase in both low-frequency (delta and theta) and high-frequency (beta and gamma) power during REM sleep is consistent with previous reports that REM is characterized by mixed-frequency EEG patterns, reflecting both high cortical arousal and periods of synchronization, which are crucial for emotional regulation and dreaming (Nir and Tononi 2010); (Insana 2020); (Llinás and Ribary 1993).

One of the most novel and surprising findings from this study is the inverse relationship between spectral power in multiple frequency bands and N3% across several sleep stages, including N1, N2, and REM. Previous research has consistently emphasized the importance of slow-wave activity (SWA), particularly delta power, as a marker of deep, restorative sleep (Tononi and Cirelli 2006)(Borbély & Achermann, 1999). However, our data suggest that reductions in spectral power across multiple frequencies, especially in sigma (12–15 Hz), beta (16–25 Hz), and gamma (25–50 Hz) ranges, are associated with higher N3%, indicating that lower brain activity during N1 and N2 may facilitate deeper sleep. This contrasts with existing studies that highlight the role of spindle activity in NREM sleep in enhancing slow-wave sleep (D.-J. Dijk 2009); (Fogel SM et al. 2007), suggesting a potential trade-off between spindle generation and the ability to enter deep sleep in some individuals.

The negative correlation between low delta power (0.1–0.5 Hz) during N2 and both N3% and total sleep duration (TST) is another surprising finding. Delta activity has long been considered essential for the restorative processes of sleep, particularly during N3 (Achermann and Borbély 1997); (Massimini et al. 2005), but its role during N2 is less understood. Our results suggest that excessive low-frequency power in N2 may hinder deeper sleep stages, potentially disrupting the balance between early-stage sleep and subsequent slow-wave sleep. This is consistent with previous discussions on the potential for excessive SWA during lighter NREM stages to disrupt overall sleep architecture (Willi et al. 1998), challenging the traditional view that more delta activity is always beneficial. This novel finding adds nuance to the understanding of how different levels of delta power during early NREM stages affect transitions into deeper sleep.

The correlations between REM percentage and spectral power across different stages provide further insight. The weak but significant negative correlations between high-frequency activity (alpha to gamma) during NREM stages and REM% align with studies suggesting that excessive cortical activation can interfere with REM sleep (Bódizs et al., 2005). More interestingly, our finding that delta power during wakefulness positively correlates with REM% suggests that pre-sleep cortical downregulation may play an important role in promoting REM sleep. This observation adds a new dimension to our understanding of how pre-sleep brain states can influence subsequent sleep architecture (Borbély 1982). While much research has focused on slow-wave sleep, these findings indicate that pre-sleep brain activity, particularly in the delta band, may also modulate REM sleep regulation, a relatively underexplored area in sleep science.

### Correlations Between Temporally Regressed Spectral Power and Sleep Metrics

The temporally regressed spectral power analysis provided even more detailed insights into the dynamic relationships between EEG spectral activity and sleep metrics. A key finding from this analysis is the strong positive correlation between N3% and increases in delta power over time during N2, indicating that the buildup of slow-wave activity over the night is crucial for deep sleep. This supports the widely accepted theory of sleep homeostasis, in which slow-wave activity reflects the brain’s recovery needs and increases in response to sleep pressure (Borbély 1982); (Riedner et al. 2007). Similarly, the negative correlation between decreases in delta power during N3 and increases in N3% suggests that individuals who lose slow-wave activity too early might require more deep sleep overall. These results partially align with earlier work showing that the timing and maintenance of slow-wave activity are critical for the restorative function of sleep (Mander, Winer, and Walker 2017); (Massimini et al. 2004).

The results for REM% are also noteworthy. Temporal increases in delta power during the awake period positively correlated with REM%, suggesting that individuals who exhibit stronger cortical downregulation before falling asleep are more likely to experience higher amounts of REM sleep. This finding supports the idea that pre-sleep brain states play a critical role in determining sleep quality and the balance of sleep stages (Borbély et al. 1981). Moreover, the negative correlation between decreases in delta power during N3 and REM% suggests that increasing slow-wave activity over time may decrease the amount of REM, which has not been explored directly and reported in the literature.

The correlations between temporally regressed spectral power and sleep duration also provided interesting results. Temporal increases in delta power during the awake period were associated with longer total sleep time (TST), which is consistent with earlier findings showing that pre-sleep cortical downregulation is predictive of better sleep quality and longer sleep duration (Borbély et al. 1981); (D. J. Dijk et al. 1991)( Dijk et al., 1995). On the other hand, reductions in delta power over time during N2 and N3 stages were associated with more TST, reinforcing the idea that the quality of deep sleep can lead to more efficient sleep throughout the night by requiring less of it. These results provide strong evidence for the role of slow-wave activity in regulating overall sleep architecture and duration (D.-J. Dijk 2009); (Tononi and Cirelli 2006).

Finally, the correlations between temporally regressed spectral power and age highlight the age-related decline in slow-wave dynamics. Our results show that delta power in N3 decreases over time in younger individuals and increases over time or potentially remains the same in older individuals, which overlaps with previous reports that aging is associated with a disruption of optimal slow wave activity and dynamics (Mander, Winer, and Walker 2017); (Ohayon et al. 2004). The decrease in power across various frequencies (very low delta, sigma, beta, and gamma) over time during N3 as it corresponds to older age goes against reports that aging individuals exhibit greater cortical activation during sleep, which may contribute to increased sleep fragmentation and reduced sleep efficiency (Carrier et al. 2001); (Feinberg and Campbell 2010). These findings suggest that interventions aimed at sustaining slow-wave activity during N3 could help improve sleep quality in older adults, who often suffer from sleep fragmentation and shorter periods of restorative sleep (Mander, Winer, and Walker 2017).

The findings from this study have important implications for sleep research and clinical practice. First, the novel observation that lower levels of spectral power during N1 and N2 are associated with increased N3% suggests that interventions aiming to reduce high-frequency brain activity in early sleep stages could promote deeper sleep. These insights could be used to develop personalized sleep therapies targeting specific frequency bands to improve sleep architecture.

One of the most promising applications of these findings is in the development of non-invasive brain-computer interfaces (BCIs) for sleep modulation. Sensory brain stimulation seems particularly promising for these purposes. BCIs that monitor EEG spectral power in real-time could be used to detect deviations from optimal sleep architecture and deliver interventions to correct them. For example, our findings suggest that BCIs could be programmed to detect excessive sigma or beta activity during N1 or N2 and provide neurofeedback or non-invasive stimulation to promote cortical downregulation, facilitating transitions into deeper sleep. Auditory or electrical stimulation targeted to enhance slow-wave activity during sleep has been shown to improve sleep quality and memory consolidation (Ngo et al. 2013); (Marshall et al. 2006), and these technologies could be further optimized using insights from spectral power dynamics.

In addition, BCIs could be designed to provide real-time feedback during the awake period to promote cortical downregulation before sleep, thereby enhancing REM sleep and overall sleep quality. By tracking pre-sleep delta power, BCIs could deliver feedback to help individuals achieve more optimal brain states for sleep onset, improving both sleep duration and stage transitions (Borbély et al. 1981); (Riedner et al. 2007). This could be particularly useful for individuals with insomnia or other sleep disorders, where disruptions in the balance of spectral power across sleep stages are common.

Furthermore, our findings on the relationship between spectral power and age suggest that BCIs could be used as diagnostic tools for detecting early signs of sleep deterioration in aging populations. By monitoring delta and gamma power dynamics across the night, BCIs could help identify age-related declines in sleep quality and provide personalized interventions to counteract these effects, such as stimulation to prolong slow-wave activity in N3 sleep. Such interventions could mitigate the cognitive and physical declines often associated with poor sleep in older adults (Mander, Winer, and Walker 2017).

## Limitations

Several limitations of this study should be noted, particularly in relation to the generalizability of the findings and the methodologies employed. First, while the sample size of 1137 participants provides a robust dataset, there are inherent limitations in the diversity of the sample that may affect the generalizability of the results. Factors such as age, health status, and individual sleep habits were not fully controlled, which may limit the application of these findings to broader populations. For example, although age was considered, other variables like medication use, lifestyle factors, or undiagnosed conditions could have influenced the EEG patterns and the relationships observed between spectral power and sleep metrics.

Another important limitation is the cross-sectional nature of the study. By relying on single-night EEG recordings, the study provides only a snapshot of each participant’s sleep patterns. This design limits the ability to infer long-term trends or causal relationships, particularly with regard to age-related changes in sleep architecture. Longitudinal studies tracking changes in spectral power and sleep metrics over time are necessary to establish how sleep evolves and how factors like aging or lifestyle interventions might influence these dynamics.

Finally, the use of correlation analyses in this study limits the ability to draw firm conclusions about causality. While significant associations between spectral power and sleep metrics were observed, it remains unclear whether these spectral changes cause better sleep or if they are simply correlated with other underlying processes. Future studies that involve experimental manipulations, such as using neurostimulation to modulate specific frequency bands, could help determine the causal relationships between brain activity and sleep quality.

## Future Directions

To build on the findings of this study, future research should aim to integrate additional physiological and behavioral metrics alongside EEG spectral power analysis. Incorporating measures such as heart rate variability, respiration, and hormonal data, as well as cognitive performance assessments, would provide a more comprehensive understanding of how brain activity during sleep influences overall health and daytime functioning. These combined measures would allow researchers to explore the complex interactions between sleep, physiology, and behavior in greater depth.

Experimental manipulations, such as using non-invasive brain stimulation techniques like transcranial electrical stimulation (tES), deep brain magnetic stimulation (DBMS) or sensory (visual and/or auditory) brain stimulation, could help establish causal relationships between spectral power and sleep quality. These interventions would offer insights into how specific frequency bands, such as slow-wave activity or sleep spindles, contribute to sleep architecture and could be used to develop targeted therapies for improving sleep in clinical populations.

Finally, future research should expand to cross-cultural and population-based studies to examine how different environments, lifestyles, and stressors influence sleep architecture and spectral dynamics. This would provide a broader understanding of sleep variability across populations and could inform public health strategies aimed at improving sleep and preventing sleep-related health issues on a global scale.

## Conclusion

In summary,(n.d.) this study provides a detailed analysis of EEG spectral power across different sleep stages and transitions, revealing both established findings and novel insights. We confirmed known patterns such as the decrease in high-frequency power during transitions to NREM sleep and the increase in low-frequency power during deep sleep, while also identifying unexpected inverse correlations between spectral power and N3 percentage, suggesting that lower brain activity in earlier stages facilitates deeper sleep. Additionally, the relationships between spectral power and key metrics like REM percentage, sleep duration, and age offer new perspectives on how sleep quality is regulated and declines with aging. The results highlight the potential for personalized sleep interventions using brain-computer interfaces (BCIs) that monitor and modulate spectral power in real-time to optimize sleep architecture and improve overall health. These findings open the door for future research to explore causal relationships and develop targeted therapies for sleep enhancement.

## Data Availability Statement

The datasets used and analyzed in this study are available from the corresponding author upon reasonable request

## Acknowledgments

This research was supported by the National Science Foundation Small Business Innovation Research Program (NSF/SBIR) under Grant No. 2146931. The authors gratefully acknowledge the invaluable assistance of Roma Shusterman in editing and refining this manuscript, whose insights and contributions significantly improved the clarity and presentation of the work.

## Financial Disclosure

This research was conducted by NeuroLight, Inc., a commercial medical device company. All authors are employees of, consultants to, or shareholders in NeuroLight, Inc., which is actively developing products related to the research described in this manuscript. The company may choose to file related patent applications.

## Non-financial Disclosure

The authors declare no non-financial competing interests related to this research. All relationships disclosed pertain to financial interests in NeuroLight, Inc., and do not influence the integrity or impartiality of the research presented in this manuscript.

## Notes

### Competing Interest Statement

The authors declare the following financial interests/personal relationships, which may be considered as potential competing interests: This research was conducted by NeuroLight, Inc., a commercial medical device company. All authors are employees of/consultants to/shareholders in NeuroLight, Inc., which is developing products related to the research described in this paper. The company may choose to file related patent applications.

### Summary of Updates

Revised several sections of the manuscript.

